# Partially Overlapping Brain Networks for Singing and Cello Playing

**DOI:** 10.1101/257923

**Authors:** Melanie Segado, Avrum Hollinger, Joseph Thibodeau, Virginia Penhune, Robert J. Zatorre

## Abstract

This research uses an MR-Compatible cello to compare functional brain activation during singing and cello playing within the same individuals to determine the extent to which arbitrary auditory-motor associations, like those required to play the cello, co-opt functional brain networks that evolved for singing. Musical instrument playing and singing both require highly specific associations between sounds and movements. However, vocal motor control is an evolutionarily old human trait and the auditory-motor associations used for singing are also used for speech and nonspeech vocalizations. This sets it apart from the arbitrary auditory-motor associations required to play musical instruments. The pitch range of the cello is similar to that of the human voice, but cello playing is completely independent of the vocal apparatus and can therefore be used to dissociate the auditory-vocal network from that of the auditory-motor network. While in the MR-Scanner, 11 expert cellists listened to and subsequently produced individual tones either by singing or cello playing. All participants were able to sing and play the target tones in tune (within 50Cents). We found that brain activity during cello playing directly overlaps with brain activity during singing in many areas within the auditory-vocal network. These include primary motor, dorsal pre-motor, and supplementary motor cortices, the primary and periprimary auditory cortices within the superior temporal gyrus including Heschl’s gyrus, anterior insula, anterior cingulate cortex, and intraparietal sulcus, and cerebellum but, notably, exclude the periaqueductal grey and basal ganglia. Second, we found that activity within the overlapping areas is positively correlated with, and therefore likely contributing to, both singing and playing in tune determined with performance measures. Third, we found that activity in auditory areas is functionally connected with activity in dorsal motor and pre-motor areas, and that the connectivity between them is positively correlated with good performance on this task. This functional connectivity suggests that the brain areas are working together to contribute to task performance and not just coincidently active. Last, our findings showed that cello playing may directly co-opt vocal areas (including larynx area of motor cortex), especially if musical training begins before age 7.

## 2 Introduction

Playing musical instruments and singing both result in musical pitch patterns by integrating auditory perception with fine-motor control. Thus, an interesting question is whether the neural systems that control these two types of musical expression are similar. Auditory-motor integration for singing relies on neural systems for vocalization, where there is a relatively direct link between a motor action and the pitch produced. This evolutionarily old auditory-vocal system comprises auditory, motor and pre-motor regions in the dorsal stream, as well as the cerebellum, basal ganglia and brainstem structures. Fig. 1 (Kleber et al. 2014). However, the auditory-vocal system is used for both speech and non-speech sounds as well as for singing. Furthermore, the vocal motor system follows a developmental sequence and does not require explicit training, at least for production of simple songs, which children produce by imitation, as with speech (Tourville et al. 2008). Nonetheless, musical instrument playing, which does require explicit training and often thousands of hours of practice, has been shown to rely on many of the same structures in neuroimaging studies (Zatorre et al. 2007). However, no previous studies have directly compared the brain networks engaged by singing and instrument playing. This comparison would allow us to assess whether learned auditory-motor associations involved in playing an instrument build on existing brain networks that are in place for vocal production, or whether they engage different or additional systems.

**Fig 1.**
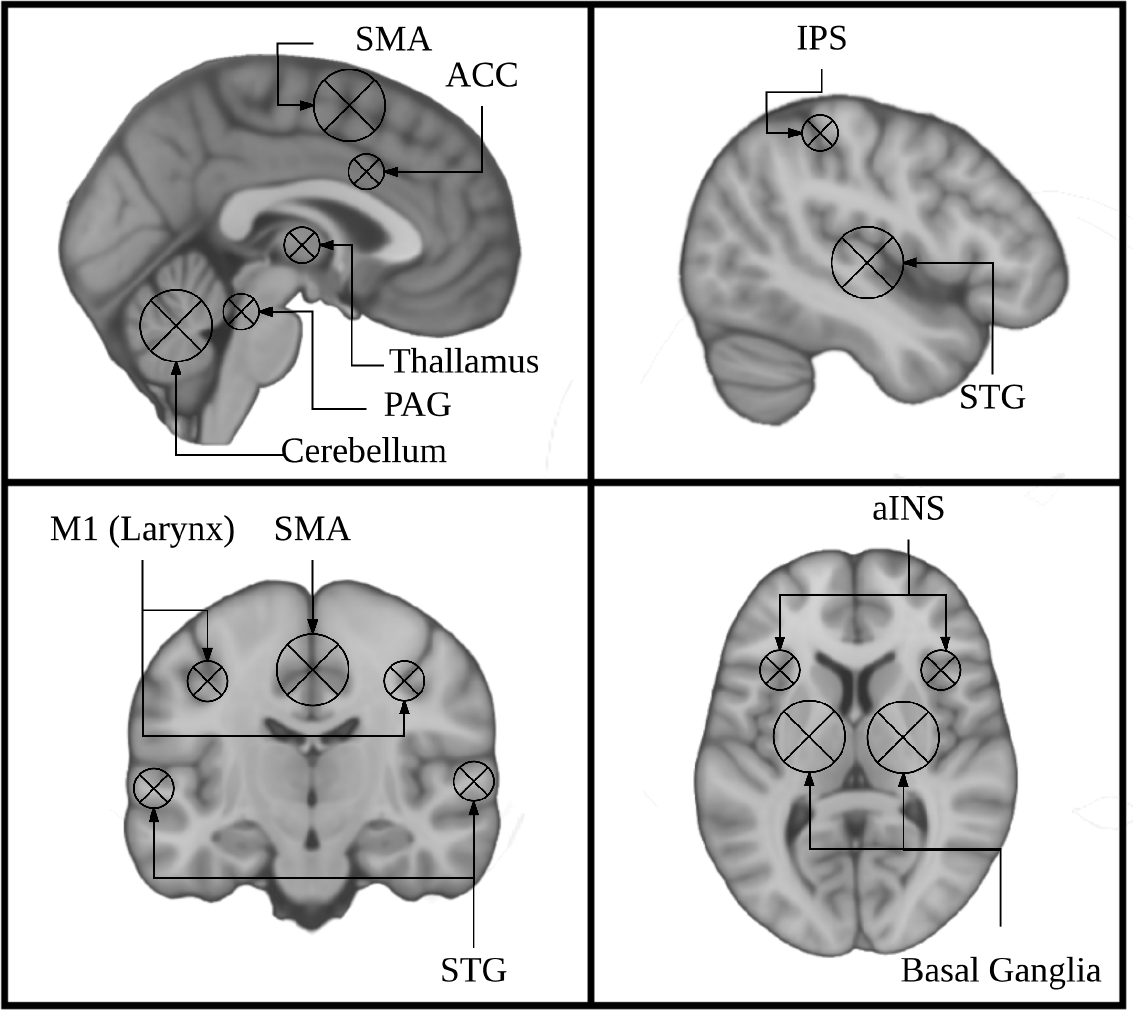
key regions identified using singing as a model instrument provide a framework for interpreting findings from research on musical instrument playing.

Current models of auditory-motor integration for music and speech comprise a feed-forward and a feed-back component (Kleber et al. 2014; Zatorre et al. 2007; Tourville et al. 2011; S. Brown et al. 2004). The feed-forward component encompasses brain areas that are responsible for motor planning and motor execution. These include primary motor, dorsal pre-motor, and supplementary motor cortices (M1, dPMC, SMA), brainstem nuclei including the periaqueductal grey (PAG), and the cerebellum. The feedback component encompasses brain areas that process sensory feedback and compare it to the expected output. Notably these include the primary and periprimary auditory cortices within the superior temporal gyrus (STG) including Heschl’s gyrus, the planum temporale, and planum polare (HG, PT, PP), as well as the superior temporal sulcus (STS). These also include the anterior insula (aINS), anterior cingulate cortex (ACC), and intraparietal sulcus (IPS). These models were informed by research on speech and non-speech vocalizations, singing, and musical instrument playing. The earliest work done on the neural correlates of vocalizations was done in non-human primates. This work showed that stimulation of the PAG induces vocalizations, and that lesioning this region leads to muteness (Jürgens 1976; Dujardin et al. 2005). Work in non-human primates has also shown that the SMA/pre-SMA and ACC are important for initiating voluntary vocalizations. (Gooler et al. 1987; Kirzinger et al. 1982). This same work helped to identify a larynx specific region of primary motor cortex in non-human primates (for review see: (Jürgens 1976). Several studies have since characterized a larynx-specific area of M1 in humans (S. Brown et al. 2008; Grabski et al. 2012) and replicated the finding that vocalization tasks recruit the PAG, ACC, and Pre-SMA/SMA in humans (Schulz et al. 2005) for review see: (Kleber et al. 2014).

Building on findings from early animal models of vocal control, research on singing has been instrumental in characterizing both behavioural features of vocal control in humans (Parlitz et al. 1999) and the associated brain regions. These brain regions comprise the auditory-vocal network. (Perry et al. 1999; S. Brown et al. 2004; Zarate et al. 2005; Kleber et al. 2007; Larson et al. 2008). Some studies have specifically investigated the feed-forward components of the auditory-vocal network by masking sensory feedback during singing (Kleber et al. 2017), asking participants to ignore perturbed auditory feedback (Zarate et al. 2008), or anesthetizing the vocal cords (Kleber et al. 2013). These studies have highlighted the role of the aINS, showing that its activity is modulated during singing when sensory feedback is masked or perturbed. Components of these studies have specifically focused on the effects of perturbed auditory feedback and found that the ACC and and IPS are involved in compensating for these perturbations (Zarate et al. 2008).

These studies also highlight the effects of musical expertise. Areas within the basal ganglia (putamen) were more active for experts than for non-experts (Zarate et al. 2008), while the aINS was less active for experts than for non-experts (Kleber et al. 2013). However, the network of areas recruited is extremely stable throughout the singing literature and even shows some degree of overlap with the areas recruited for speaking in both auditory and motor regions (Hickok et al. 2003; Ozdemir et al. 2006). Differences between these two systems, like the relative right lateralization of auditory cortex for singing compared to speech, are thought to be related to the increased dependence on pitch processing during singing (Ozdemir et al. 2006). It therefore would seem likely that musical instrument playing, which is also highly dependent on pitch processing, would show a high degree of overlap with the brain areas recruited of singing.

Functional imaging research on musical instrument performance has also helped inform our understanding of the auditory-motor integration system. Studies on piano and keyboard playing (R. Brown et al. 2015; Baumann et al. 2005; Bangert et al. 2006; Haslinger et al. 2004; Parsons et al. 2005), simulated violin performance (Lotze et al. 1999; Lotze et al. 2003), simulated guitar playing (Vogt et al. 2007; Buccino et al. 2004; Higuchi et al. 2012), and playing a trumpet (Gebel et al. 2013) show activation in many of the same core auditory and motor regions as are seen in the auditory-vocal network, but notably does not include the brainstem. However, as noted above, no direct comparison has been performed between the brain areas recruited for vocal and instrumental pitch production. To directly compare singing and instrument playing we need to use an instrument that has a continuous (as opposed to discrete) pitch mapping like the voice, but whose control is completely independent of the vocal apparatus. Instruments in the violin family fit both criteria. fMRI research on violin playing has relied on imagined performance or finger tapping as a proxy for real performance because a violin could not be played in the scanner (Lotze et al. 2003). Consequently, this research lacked the auditory feedback necessary to directly compare singing to playing. One study made use of an fMRI compatible trumpet, but trumpet playing is not entirely independent of the vocal apparatus (Gebel et al. 2013). Other fMRI research on musical instrument playing has focused on the keyboard (R. Brown et al. 2015; Chen et al. 2012). However, the keyboard differs from the voice in two key ways. First, the keyboard is a discrete pitch instrument, in which the pitch of each note is mapped to a key in a one-to-one manner, and cannot be changed once a key has been pressed. In contrast, the human voice is a continuous pitch instrument which can produce any pitch within a range, not just discrete values, and can change this pitch on an ongoing basis (notably, in response to a feedback perturbation) (Parlitz et al. 1999; Burnett et al. 1998). This continuous nature of pitch is paralleled in fretless string instruments from the violin family. Second, in keyboard playing, pitch and timing are controlled by the same movement, a key press. In singing, pitch and timing are often decoupled; pitch is controlled by the tension of the vocal cords sound onsets and offsets are controlled primarily by the diaphragm. This distinction is paralleled by the asynchronous movements of the left and right hands required to play instruments from the violin family, which control pitch value and timing of sound onsets and offsets, respectively.

In the present study we directly test the hypothesis that singing and cello playing recruit an overlapping network of brain regions in auditory, motor, and auditory-motor integration regions. Specifically, we are interested in the auditory-vocal network Fig. 1. To accomplish our goal, we have developed an MR compatible cello device where sound feedback is delivered directly to the player during scanning (Hollinger et al. 2013; Hollinger et al. 2015) Fig. 2. This cello uses optical sensors embedded in the fingerboard and bridge to capture finger position and string vibration, respectively. The optical sensor on the bridge of the cello provides realtime, analog sound feedback. The MR cello lacks a resonant body, but the length of the fingerboard and strings, and, by extension, the locations of where fingers would be placed to produce specific notes are the same as those found on a standard full-size cello. We also constructed a miniaturized bow to fully leverage the continuous pitch nature of the cello. By comparing the neural correlates of singing to those of cello playing directly in the same individuals, we can determine the extent to which musical instrument playing makes use of functional brain networks that have evolved for singing. We expect that cello playing and singing will recruit largely overlapping areas in premotor and supplementary motor areas, auditory cortices, and the cerebellum. We also expect that activation in primary motor cortex will be specific to the hand and larynx areas, respectively. Additionally, because playing in tune requires auditory-motor integration, we expect to see activation in areas that are thought to be responsible for perceptual monitoring and error correction (IPS, ACC, aINS and thalamus). We also expect to see functional connectivity between these regions during both singing and cello playing, reflecting the network properties of these regions acting in concert to accomplish the task.

**Fig 2.**
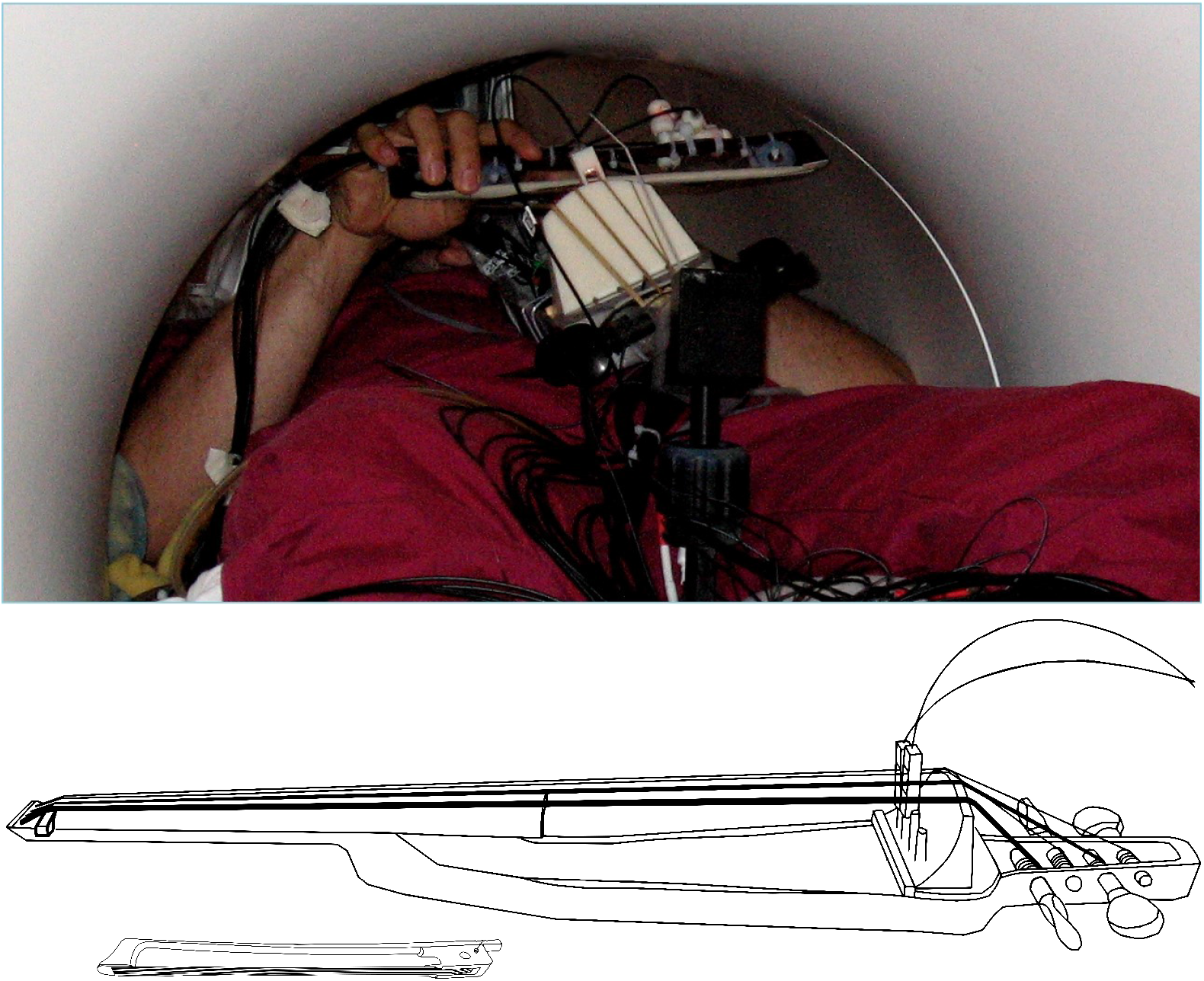
Top: View of the MR-compatible cello and bow inside the scanner during experiment. Bottom: Line drawing of cello and bow developed for use inside the MR Scanner. Cello fingerboard has the same dimensions as a fullsized cello to preserve the location and spacing of notes. Optic sensors at the bridge are used to detect string vibrations, and are converted to analog audio output by a custom ARM board (not shown). The MR Compatible bow is 20cm long, which is just under a third of the length of a typical cello bow (apx.70cm)

## 3 Materials and Methods

### 3.1 Subjects

A total of 12 expert cellists (6 Female) were recruited from the Montreal community. All participants were right handed, had normal hearing, did not report any neurological disorders, and had no contraindications for the MRI environment (mean age = 21.4, mean years experience = 13.9, mean starting age = 7.4, mean practice hours per week = 24.8). Eleven participants were included in the final analysis, one participant was excluded from the analysis due to an equipment failure. This study was approved by and carried out in accordance with the recommendations of Montreal Neurological Institute Research Ethics Board and the McConnell Brain Imaging Centre. All subjects gave written informed consent in accordance with the Declaration of Helsinki.

### 3.2 Experimental paradigm

#### 3.2.1 Stimuli and task conditions

Participants were asked to listen to and subsequently produce target tones both singing and on an MR-Compatible cello. There were three experimental conditions: Sing/Play, Listen, and Rest. For all three conditions, we presented auditory target tones that were two seconds long. Presented tones were either E3, F^#^3, G^#^3 for cello, and E3, F^#^3, G^#^3 or E4, F^#^4, G^#^4 for singing depending on participant’s vocal range. Tones were recorded by either a female vocalist, male vocalist, or on a cello. Tone presentation was followed by a pre-recorded auditory instruction to either Listen, Sing/Play, or Rest. On some trials auditory feedback was either masked or pitch shifted (see below).

Participants underwent a familiarization session followed by an fMRI session. For both the familiarization session and the fMRI session, a microphone was suspended approximately 2 inches from their mouth, the MRI compatible cello device was placed along their torso using a specialized support, and headphones were provided (Sensimetrics S14 fMRI insert headphones, Dayton Audio DTA-1 amplifier). The microphone and cello were connected to a Mackie 802VLZ4, 8-channel mixer and to a midi controlled TCHelicon VoiceOne pitch shifter which was used to prevent the audio feedback from reaching the headphones on certain trials. Pink noise was played through the headphones to reduce bone conduction so that audio was being delivered exclusively through the headphones. All volume levels were adjusted on a per subject basis, but on average pink noise was presented at 78.3 dB SPL A and auditory targets were presented at approximately 15.6 dB above the noise floor. The experiment was run using custom scripts written in python. All vocalizations and cello sounds were digitally recorded using a Sound Devices 744T digital recorder Fig. 4.

**Fig 3.**
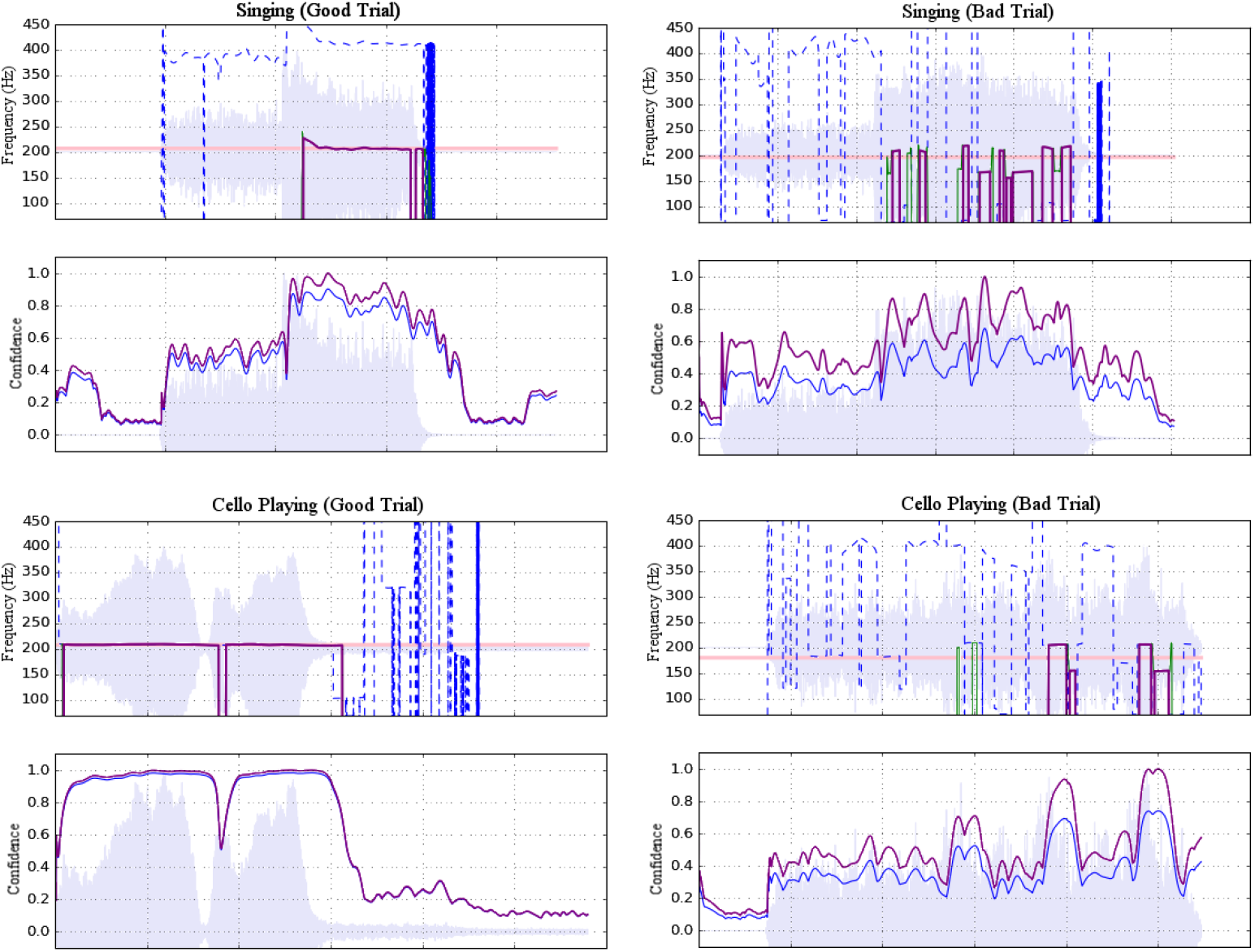
Examples of accepted (left) and rejected (right) trials for both singing (top) and cello playing (bottom). Each example shows a plot of pitch estimations above a plot of confidence ratings. For the former, dashed blue lines represent raw pitch estimation whereas solid purple lines represent potentially stable pitch estimates adjusted to the fundamental. For the latter, raw confidence ratings are shown in blue and normalized confidence ratings are shown in purple. Shading indicates amplitude envelope

**Fig 4.**
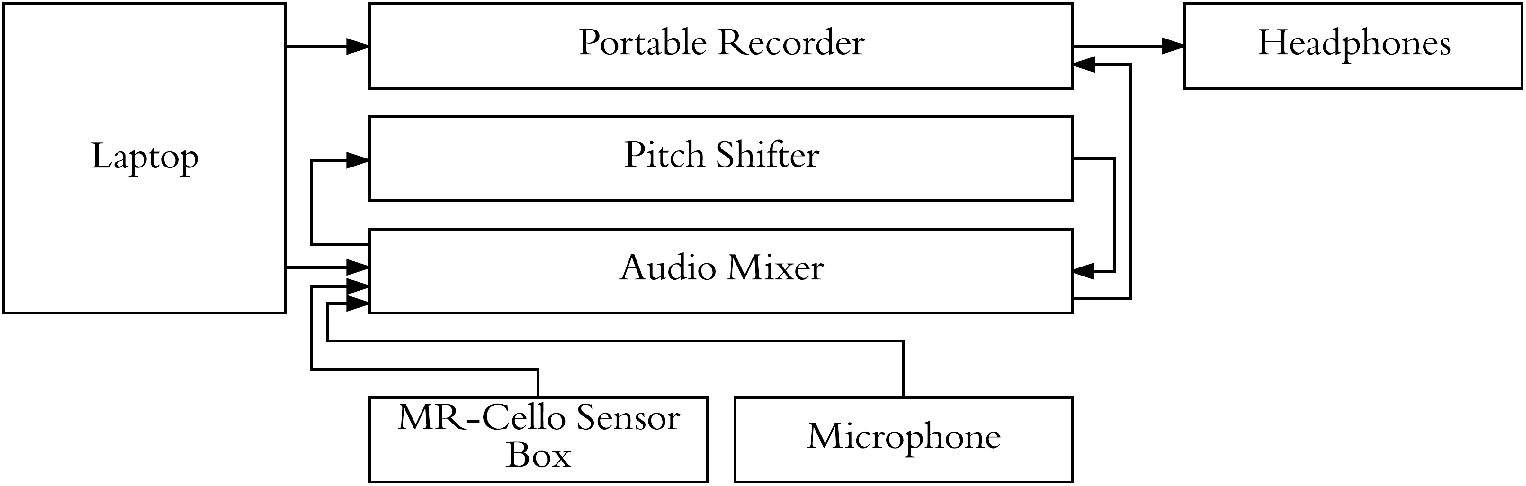
Experimental setup used to present stimuli, cello audio, and singing audio through headphones. Pitch shifter allowed for audio feedback from cello/singing to be blocked on specific trials while still presenting audio stimuli.

A sparse sampling paradigm was used for the fMRI session (Belin et al. 1999), where a long delay in TR was used to allow tasks to be carried out in the relatively silent period between functional volume acquisitions, thus minimizing acoustical interference and also avoiding movement-related artifacts since the scanning takes place after the motor production for each trial Fig. 5.

**Fig 5.**
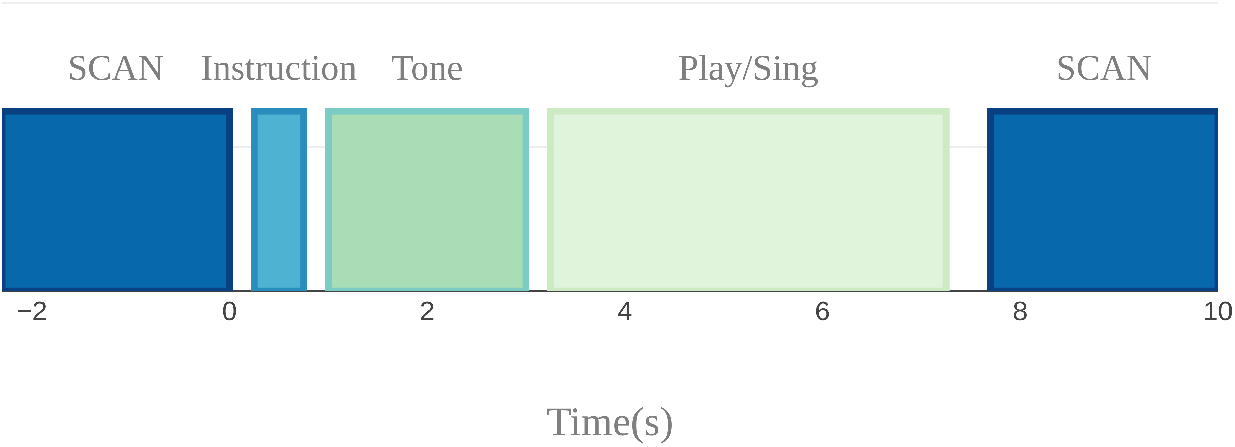
Sparse sampling design used to avoid auditory and motion artifacts. Auditory target presentation, instruction presentation, and singing and cello playing were done in the silent 7.7s period between scans

#### 3.2.2 Procedure

To allow participants to adjust to the fMRI-compatible cello and the constraints of playing it in the scanner, each person underwent a 45 minute familiarization session no more than one week prior to their session in the MR-scanner. During the familiarization session each participant was asked to lie on a foam mat inside a structure that simulated the space constraints of the MRI environment. All participants were asked to perform a series of scales, which contained all of the target notes that would be presented during the experiment and a reduced duration (10-minute version) of the experimental task.

On performance trials, participants were instructed to sing or play back the target tone for 4seconds. For singing trials, participants were instructed to sing with closed lips in order to reduce breathing artifacts in the recorded signal and movement artifacts in the fMRI signal. For cello playing trials they were instructed to use as few bows as possible to reduce movement artifacts (apx 1second per bow). Between trials participants were instructed to keep their hands on the cello and to move as little as possible. During familiarization, participants went through 2 reduced-length experimental runs (one cello, one singing), with all conditions included in each run). Three participants underwent a second familiarization session due to equipment problems during their first scheduled session.

Within one week of the familiarization session, participants were tested in the Siemens Trio 3T magnetic resonance (MR) scanner at the Brain Imaging Center of the Montreal Neurological Institute. Each participant was fitted with MR-compatible headphones. The MR-compatible microphone was attached to the mirror support system. The MR-compatible cello was laid across the torso using a special MR-compatible stand. Sound levels were adjusted on a per participant basis so that, during trials with masked auditory feedback, participants could not hear their voice or the sound of the cello above the pink noise.

During the fMRI session, participants performed two cello playing runs and two singing runs. Run order was counterbalanced across participants. During each of these runs, trial order was pseudo-randomized. Following the presentation of each target tone, participants were instructed to sing or play the cello, or to listen. On rest trials no auditory target was presented. On some of the performance trials, auditory feedback was pitch shifted (40 trials, up to one semitone) and on others it was fully masked (20 trials). Due to technical issues with the pitch shifter during data acquisition, these data were not included in the final analysis. The no audio condition and listen condition served as controls for the auditory and motor aspects of the performance trials.

#### 3.2.3 MRI acquisition

A high resolution T1-weighted anatomical scan (voxel=1mm^3^) was collected between runs 2 and 3. During the 4 functional runs, one whole-head frame of 28 contiguous T2*-weighted images were acquired (Slice order = Interleaved, TE=85ms, TR=10s, Delay in TR = 7.7s, 64×64 matrix, voxel size = 4mm^3^). All tasks were performed during the 7.7s silent period between functional volume acquisitions. As such, the tasks were done in silence. Timing of the auditory stimulus presentation was varied randomly by up to 500ms to increase the likelihood of obtaining the peak of the haemodynamic response for each task. Within each run, each condition was presented 10 times for a total of 20 acquisitions per condition for singing and 20 acquisitions per condition for cello playing. A high resolution whole brain T1-weighted anatomical scan (voxel=1mm^3^) was collected between runs 2 and 3.

### 3.3 Behavioral analyses

Individual trials of singing and cello playing were analyzed with pitch information extracted from audio signals. Each trial’s audio was first segmented from the MR-compatible microphone recording by hand using Audacity software. The trials were then processed using a custom analysis pipeline implemented in Python, with a GUI for visualizing and optimizing analysis parameters.

The ambient noise in the scanner room had a peak resonance of 160Hz that interfered with the extraction of fundamental target pitches, so harmonics 3-10 of the cello and singing tones were used for pitch extraction. To reject room noise and to isolate harmonics of interest, the raw microphone signal was high-pass filtered with a cutoff at 367Hz and low-pass filtered with a cutoff at 4216Hz. Pitch estimation was then performed using the YinFFT algorithm provided in the Python module Aubio (Brossier 2007), producing a time-series of pitch estimations (detected harmonic, in Hz) and confidence ratings (between 0 and 1). Estimates were adjusted to their representative fundamental pitches before selecting stable pitch regions for further analysis. Stable pitch regions were defined as: segments of at least 150ms in which the rate of change of the pitch did not exceed 100Hz/sec (or apx. 0.07Hz per 32-sample pitch estimation window at the sampling rate of 44100Hz). Of these regions, only those that maintained a confidence rating of at least 0.7 were included. Trials were rejected if no regions were found to meet the stability and confidence criteria (see Fig. 3). In total, 92.4% of trials were retained. Rejected trials were excluded from the fMRI analysis.

Pitch accuracy was calculated on a per-trial basis as the deviation between the produced and target pitches, expressed in cents using equation 1. An overall accuracy score for each participant was then determined by calculating their mean pitch accuracy across all trials. Finally, scores were analyzed using a two way (instrument by target) ANOVA implemented in R (R Development Core Team 2008).

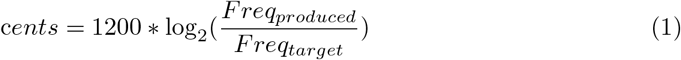

### 3.4 fMRI Analyses

fMRI data were analysed using the FSL5.0 FEAT toolbox (Jenkinson et al. 2012). Brain extraction was carried out using BET2. Functional volumes were aligned to the high resolution anatomical and then to MNI152 standard space using FLIRT linear registration with 12DOF. Motion parameters were estimated using MCFLIRT and. fMRI time course was temporally filtered to remove drifts greater than 300ms. To boost the signal to noise ratio, images were spatially smoothed with an 8mm FWHM kernel. FLAME-1 mixed effects modeling was used to fit the GLM to the fMRI signal.

Significance was determined using an FSL cluster probability threshold of p < 0.05 with a voxel-wise significance level of z=3.3 (p < 0.01). The cluster probability threshold serves as a correction for multiple comparisons. Four contrasts were carried out: Singing vs Rest, Cello playing vs Rest, Cello Playing vs Singing, and Singing vs Cello Playing. Additionally, task performance (on a per-subject basis) was regressed against the BOLD signal for each of these contrasts. Statistical conjunctions were carried out using the conjunction script created by the Warwick University Department of Statistics, which also made use of the FSL tools (Nichols et al. 2005). This script carries out a voxel-wise thresholding of p < 0.05 in both conditions of interest, and then carries out a cluster correction of p < 0.05.

Functional Connectivity analyses were carried out using the FSL5.0 FEAT toolbox. A seed region in auditory cortex was identified by masking the conjunction of singing and cello playing from the functional data with an anatomically defined mask of Heschl’s Gyrus (Harvard Structural Brain Atlas). Seed regions in primary motor cortex (M1) were identified by masking singing and cello playing with an anatomically defined mask of post-central gyrus (Harvard Structural Brain Atlas). An additional seed region was identified using a functionally defined mask of larynx area in M1 from Kleber et al. The activation time course in each of these regions was extracted and correlated with the whole-brain time course for each task of interest, which was estimated using the GLM. Correlated voxels were thresholded as described above. Regions that showed a correlated time course were then linearly regressed with task performance to determine whether those areas were contributing directly to good intonation.

## 4 Results

### 4.1 Task Performance

We first carried out a behavioural analysis to confirm that our participants, expert cello players, could sing 3 target tones and also play them on the MR-Compatible cello with at least quarter tone accuracy (50 Cents). By performing a two way anova (instrument by target tone), we found that participants could produce each of the three target tones within the specified accuracy both when singing (mean deviation from target= −8.9 Cents, stdev = 58.9 Cents) and when playing the cello (mean = −7.79 Cents, stdev = 85.39 Cents) Fig. 6. There was no significant effect of instrument (p<0.65,F = 0.21), but there was a significant effect of tone (p<7.14 × 10^−4^, F = 11.54) and a significant tone by instrument interaction (p<4.21 × 10^−8^, F = 30.65). Post-hoc tests showed that, when playing the cello, participants tended to be flat on the highest tone (mean = −39 Cents) and that, in singing, they tended to be flat on the lowest note (mean = −15 Cents). The undershoot on the highest note for cello playing was likely because the note was the most difficult to reach within the confines of the scanner. The undershoot on the lowest note for singing, while significant, is within an eighth tone of the target pitch which is well below the quarter tone threshold for considering a note in tune. There was no within-subject correlation between performance on the cello and performance on the singing trials.

**Fig 6.**
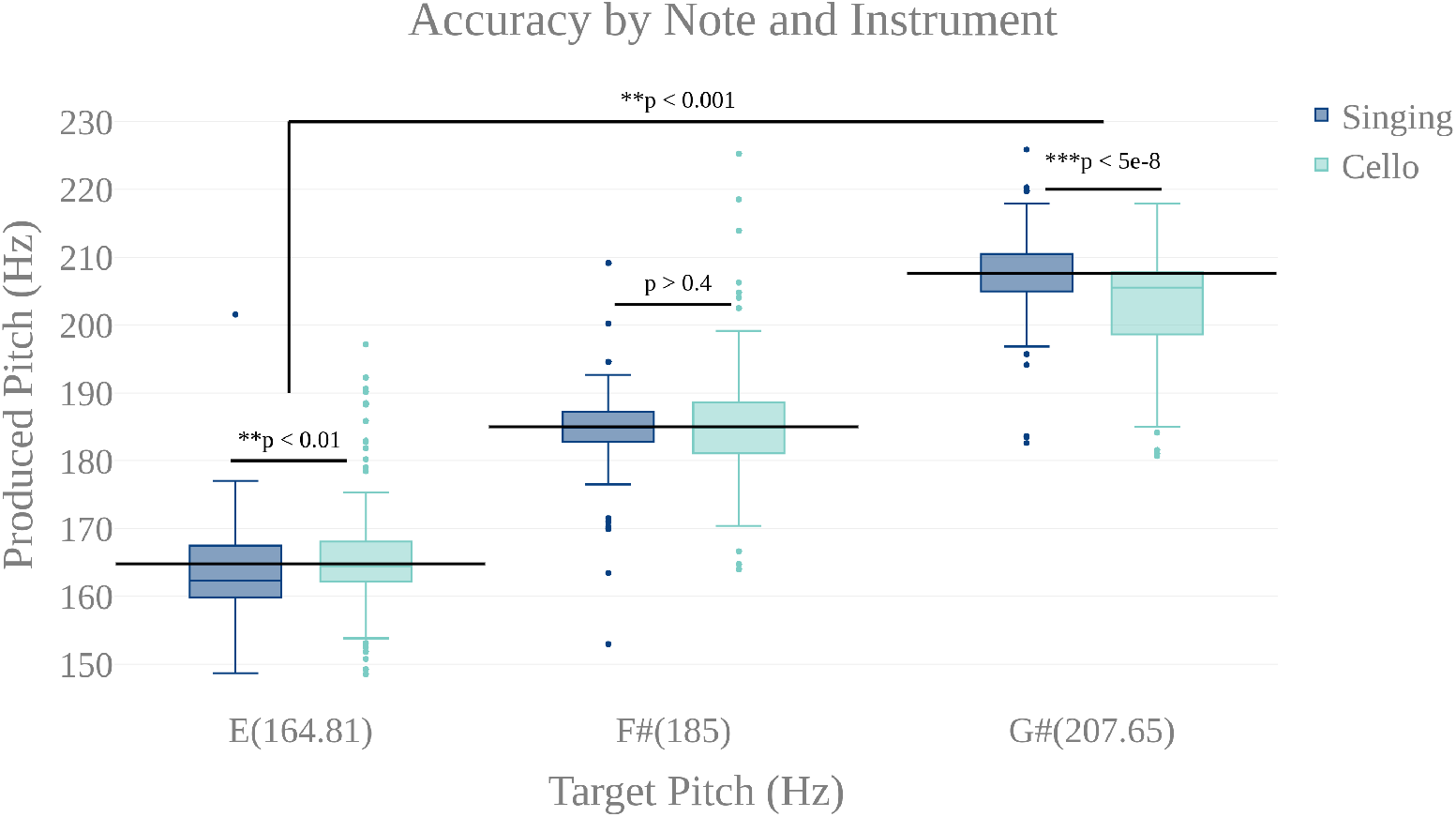
Accuracy by note for singing(dark blue) and cello playing(light blue). There is a highly significant main effect of note, but no main effect of instrument. There is a significant note by instrument interaction showing that participants were slightly flat on the lowest note when singing and slightly flat on the highest when playing the cello. However, mean produced tones were within a quarter tone accuracy(50cents)

### 4.2 fMRI Findings

#### 4.2.1 Similarities between singing and cello playing

To test the hypothesis that singing and cello playing activate a shared set of brain areas, we first identified the respective networks by performing two contrasts: Singing vs rest and Cello playing vs Rest. The Singing vs rest contrast was used to test the hypothesis that the singing task would activate the auditory-vocal network identified in previous literature (Kleber et al. 2014). We found that, consistent with the areas reported in the literature, singing activated Pre and Post central gyrus (R > L), SMA extending to ACC (bilateral), IPS (R), the length of the STG extending to supramarginal gyrus, STS, aINS (bilateral), cerebellum VI, VIIa, and CrusI-II (bilateral), thalamus extending into caudate (bilateral), the globus pallidus (bilateral), putamen (bilateral), and PAG, and the pons. In addition to areas within the auditory-vocal network, we also saw activity in middle frontal gyrus (right) Fig. 9. This contrast did not show larynx-specific activation in M1; however, we found that larynx activation was present at an uncorrected threshold of p<0.05. To confirm this finding with a more focused though statistically stringent approach, we performed a fixed-effects analysis consistent with what has been done in previous studies of larynx function (S. Brown et al. 2008). This analysis showed significant activation in larynx area of both right and left M1 (zstat_right_ = 5.3, zstat_left_ = 4.4), in locations consistent with that reported by Brown et al in M1 for phonation (d_right_ = 9.38mm, d_left_ = 9.16mm) (S. Brown et al. 2008) Eqn.2.

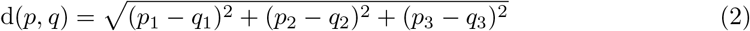

For cello playing vs rest, many of the same auditory-vocal areas were active compared to singing Fig. 9. Again, activity clusters are seen throughout the pre- and post-central gyri (bilateral), SMA extending into ACC, and IPS. Activity also extends the length of the STG and superiorly into the supramarginal gyrus. Clusters are also seen in middle frontal gyrus (right), and in the anterior insula (bilateral). In the cerebellum, activation extends through VI, VIIIa, VIIIb, VIIb, CrusI-II, Vermis CrusII, and Vermis VI. Clusters of activity are also seen in thalamus (bilateral) extending into caudate (right), and in the globus pallidus (right).

To specifically test the hypothesis that singing and cello playing both engage some of the same components of the auditory-vocal network, we carried out a statistical conjunction of the cello vs rest and singing vs rest conditions (Nichols et al. 2005). The conjunction showed overlapping activation throughout the auditory-vocal network. However, no overlap was seen in Putamen, or Brainstem. In the cerebellum, overlapping activity was seen in VI, CrusI-II, VIIIa, and VIIIb Fig. 9

Task accuracy was then linearly regressed against both singing vs rest and cello playing vs rest to determine which of the active areas were correlated with, and therefore likely to be contributing to, task accuracy. In both cases, areas that were active for the task were positively correlated with task performance (no negative correlations were observed). Specifically, for singing, all regions of the auditory-vocal network were more active as participants performed better. For cello playing, pre- and post-central gyri, STG extending into the supramarginal gyrus, aINS, cerebellum, and thalamus were more active in participants that performed better. Fig. 9

We further hypothesized that singing and cello playing would show overlap in areas whose activity contributed to higher accuracy. To test this hypothesis, we carried out a statistical conjunction of the singing vs rest and cello vs rest regressions by task performance. This conjunction showed overlap in SMA extending into ACC, IPS, middle frontal gyrus (right), STG, supramarginal gyrus, the aINS, and thalamus extending to caudate. In the cerebellum, there was overlapping activation in VIIb, VIIIa, VIIIb, and VI Fig. 9

The previous analyses found that singing and cello playing activate a shared set of brain regions, and that activity in many of these regions is positively correlated with task accuracy. Building on these findings, we decided to test the hypothesis that areas within this set are functionally connected in both singing and cello playing Fig. 10. To accomplish this goal we performed a functional connectivity analysis using the activity in Heschl’s gyrus (bilateral, from Harvard Brain Structural Atlas) as a seed region and correlating activity within this seed with activity in the rest of the brain on a voxelwise basis. For singing, we saw correlated activity in auditory cortices of the STG (bilateral) both within and around the seed area, supramarginal gyrus (bilateral), pre-central gyrus (right), inferior frontal gyrus (bilateral), and in VIIIa of the cerebellum (left). For cello playing, we saw correlated activation within the seed region and also in posterior STG extending into the supramarginal gyrus, pre- and post-central gyrus, and VIIIa, VIIIb, VIIb, VI (left), and Vermis VI of the cerebellum. The conjunction of singing and cello playing functional connectivity analyses showed shared connectivity in Heschl’s gyrus, pre-central gyrus, and VIIIa of the cerebellum (left) Fig. 10

Functional connectivity in both singing and cello playing using the same seed region was then linearly regressed with task performance to determine the correlation between connectivity and pitch accuracy. In singing, higher functional connectivity was positively correlated with performance in the supramarginal gyrus, and in cerebellar VIIa (left) and VIIb (right). In cello playing, higher functional connectivity was positively correlated with performance in pre- and post-central gyrus, the supramarginal gyrus, and cerebellar VIIIa and VI (bilateral) Fig. 10

#### 4.2.2 Differences between singing and cello playing

In order to characterize the differences in brain activity between singing and cello playing, two contrasts were first carried out: singing vs cello playing and cello playing vs singing Fig. 9. For singing vs cello playing, no significant clusters of activation were observed after correcting for multiple comparisons. However, the PAG and Putamen were active in the Singing-Rest contrast, and not in Cello Playing-Rest, or the conjunction of Cello-Rest and Singing-Rest. Additionally, at an uncorrected threshold of z=1.8 differences can be seen in larynx area of motor cortex. In order to directly test the hypothesis that the larynx area of motor cortex was more active for singing than for cello playing, which was one of our predictions, we performed a region of interest analysis of post central gyrus using the mask of laryngial motor area described above. This analysis showed more activation for singing than for cello playing in larynx area. We further tested this hypothesis by performing a 2 factor anova (Instrument by Region of Interest) with spherical ROI in hand motor regions (bilateral) and larynx area of motor cortex (bilateral). This analysis showed significant effects of both Instrument (p < 8.28 × 10^−5^, F=19.68) and ROI (p < 2.42 × 10^−2^, F=5.53), as well as a significant Instrument x ROI interaction (p < 1.03 × 10^−6^, F=34.47) driven by a highly significant difference between cello playing and singing in hand motor regions (p < 1.71 × 10^−5^, F=33.6). However, no significant difference was observed in larynx area (p < 0.13, F=2.45) Fig. 8. To understand why the contrast of singing vs cello playing showed sub-threshold differences in larynx area but the anova did not, we decided to consider potential covariates. Based on previous research showing that early trained musicians show advantages in pitch tasks, we hypothesized that early trained musicians may use larynx area of motor cortex to a greater extent than late trained musicians. To test this, we included starting age as a covariate. While starting age itself did not significantly correlate with larynx activation in either singing(r = −0.31, p < 0.34) or cello playing(r = −0.41, p < 0.26), the contrast of singing vs. cello playing in larynx area was significant when accounting for the effect of starting age (p < 0.0091, F=8.3) Fig. 7. Additionally, when data were divided into early and late starting groups (as in (Penhune 2011), cellists that began musical training before age 7 (n=5) showed significantly more activation in larynx area during cello playing than did those that started age 7 and up (n=6) (p<0.01, F=10.26) Fig. 7.

**Fig 7.**
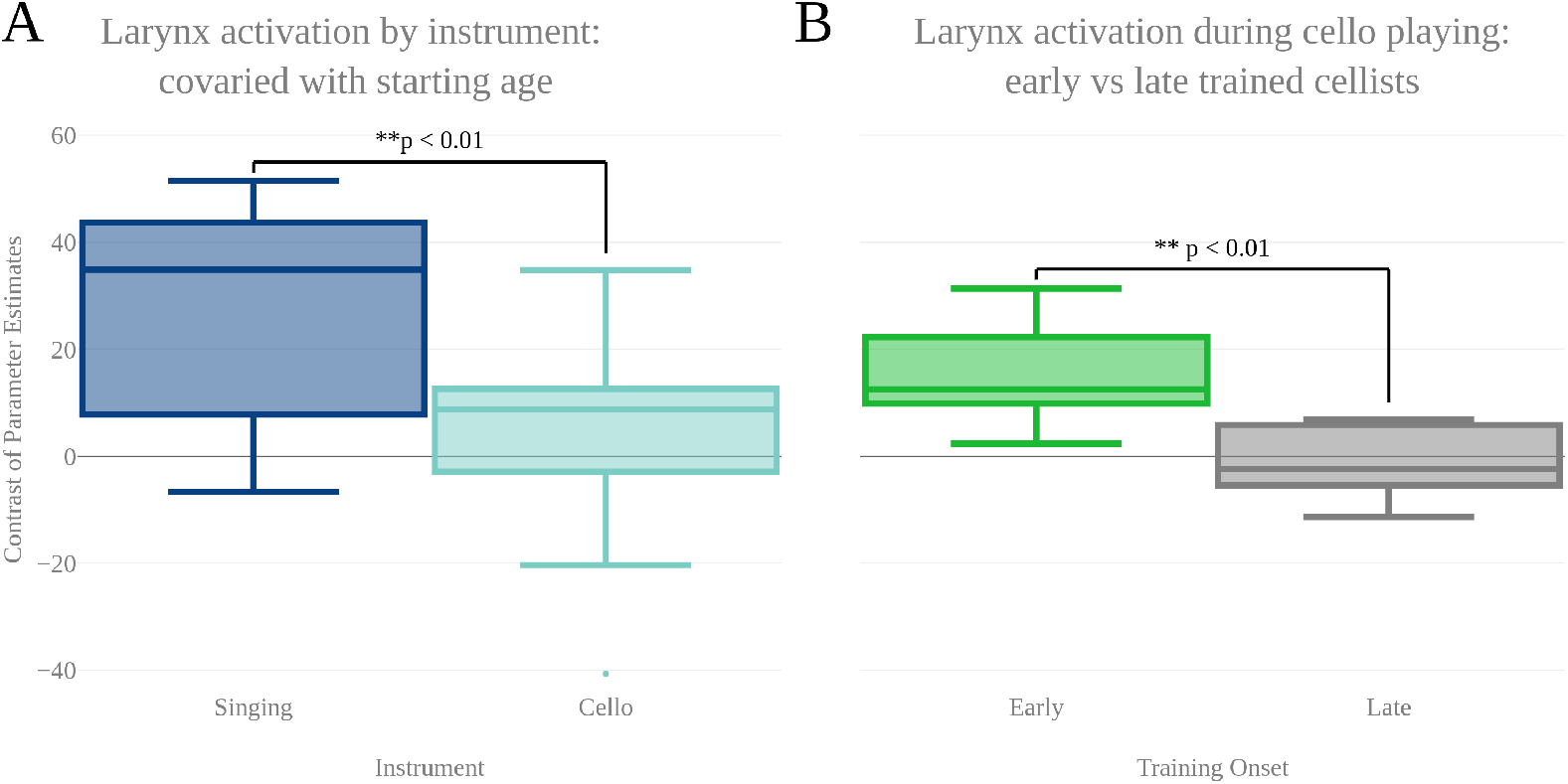
A: Contrast of parameter estimates (units arbitrary) in larynx area of motor cortex for singing (blue) and cello playing (turquoise). When starting age is included as a covariate, anova shows a significantly more activity in larynx area during singing than cello playing. B: Mean contrast of parameter estimate values in larynx area of motor cortex during cello playing for Early trained (green) and Late trained (grey) cellists. Cellist that started training before age 7 show significantly more activity in larynx during cello playing than do those that started age 7 and later

**Fig 8.**
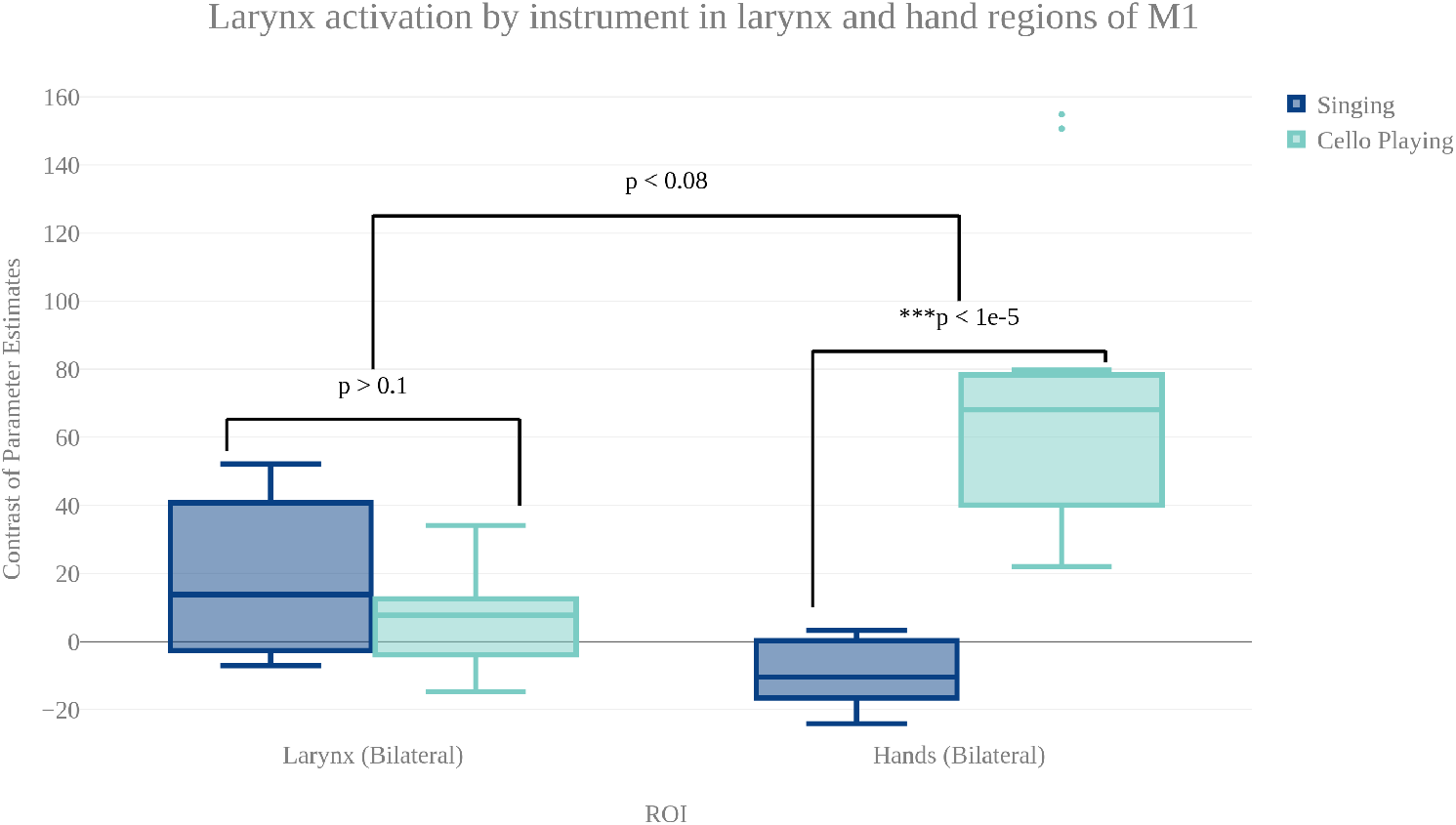
Contrast of parameter estimates (units arbitrary) in larynx and hand area of motor cortex for singing (blue) and cello playing (turquoise). The anova showed no significant effect of instrument or brain area, but did show a significant instrument by region interaction.

For cello playing vs singing, there was more activation in STG extending to the SMG, pre- and post-central gyri, and in the posterior insula. In the cerebellum, cello playing showed more activation than singing in I-IV, in VI and in VIIIb (bilateral). In pre- and post-central gyri as well as in the cerebellum, the activation peaks were centered on areas associated with hands/arms (Yousry et al. 1997).

No significant differences in functional connectivity were observed when directly contrasting singing and cello playing.

## Discussion

### 4.3 Findings

Four conclusions may be reached from this research study. First, that brain activity during cello playing directly overlaps with brain activity during singing in many areas within the auditory-vocal network. These areas include dorsal motor and premotor areas, SMA and Pre-SMA, STG, ACC, aINS, IPS(R), and Cerebellum but, notably, exclude the PAG and Basal Ganglia (Putamen). Second, that activity within the overlapping areas is positively correlated with, and therefore likely contributing to, both singing and playing in tune as shown by correlations with performance measures. Third, that activity in auditory areas is functionally connected with activity in dorsal motor and pre-motor areas, and that the connectivity between them is positively correlated with good performance on this task. This functional connectivity suggests that the brain areas are working together to contribute to task performance and not just coincidently active. Last, our findings showed that cello playing may directly co-opt vocal areas (including larynx area of motor cortex), especially if training begins before age 7.

### 4.4 Questions Answered

#### 4.4.1 Neural Correlates of Learned, Arbitrary Associations

This study provides evidence that relatively new auditory-motor integration tasks like stringed instrument playing make use of the auditory-vocal network, which is thought to be an evolutionarily old system Fig. 9. The interpretation that cello playing makes use of neural mechanisms that evolved for singing is consistent with the theory of Neuronal Recycling proposed by Dehaene et al (Dehaene et al. 2007). This theory proposes that cultural tasks (like arithmetic) are too new to be the product of evolution and that, as a result, they have to make use of cognitive mechanisms that are in place for more basic tasks (like direction processing). We propose that our findings are an example of the same concept but in the auditory-motor domain. The auditory-vocal network used for singing develops without explicit training, much like spatial processing in the visual domain. After explicit instruction, cello playing brain activity patterns overlap with singing throughout the auditory-vocal network. Potentially the best point of evidence supporting this interpretation is our finding that cellists that began training before age 7 playing activated the larynx area of motor cortex during cello playing despite cello playing not relying on the larynx. We cannot rule out the possibility that cellists were humming subvocally during the cello playing task, though we did rule out the possibility that they were actually humming using the continuous microphone recording. However, if subvocal humming was responsible for the larynx activation observed during cello playing, it seems likely that other vocalization specific areas like the PAG would also be recruited, which is not the case, and seems unlikely that we would see a starting age effect. In addition, no significant activation was observed in basal ganglia or brainstem areas that, in singing, are active even during imagined singing (Kleber et al. 2007).

**Fig 9.**
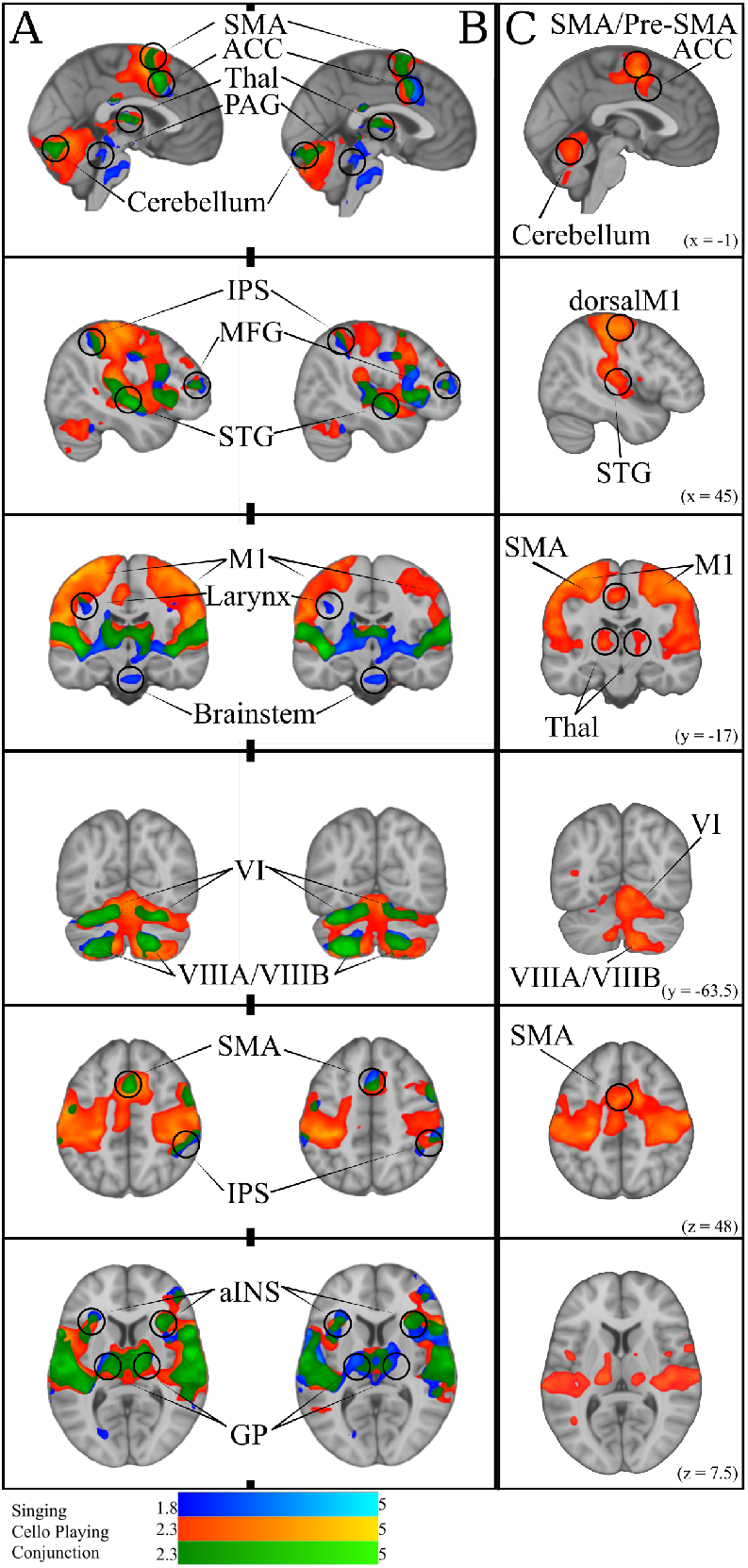
A: Simple Singing (blue), cello playing (orange), and conjunction (green). B: Simple singing (blue), cello playing (orange) and conjunction (green) regressed by task accuracy with better performance correlated with more activation. The conjunction of singing and cello playing shows overlapping activation in SMA, ACC, Thalamus, PAG, IPS, MFG, STG, MI (larynx area for singing), aINS, GP, and VI, and VIIIA/VIIIB of the cerebellum. All of these regions were positively correlated with better intonation. C: Cello playing > Singing throughout dorsal motor and premotor regions, SMA/Pre-SMA., posterior STG, Thalamus, and Cerebellum

#### 4.4.2 The Role of the ACC and aINS

Some brain areas, like the ACC and aINS, are reported to be active both in studies of musical instrument performance and singing, but are given different interpretations according to the task. In studies of piano performance, the activity of the ACC is thought to reflect coordination of hand movements (Parsons et al. 2005) while in studies of singing the ACC is thought to be specifically involved in initiating vocalizations (Kleber et al. 2007) similar to previous work done in animal models (Jürgens 2002). Similarly, the aINS is thought to integrate somatic information from the body to support bi-manual coordination in piano playing (Parsons et al. 2005) and to coordinate the vocal musculature during singing (Kleber et al. 2007) together with other interoceptive inputs (Kleber et al. 2013). In our study, we found that activity in the aINS and ACC was directly overlapping during cello playing and singing. Consequently, we propose that the role of the aINS and ACC is to coordinate the movements of whichever motor effectors are required in order to produce pitched sounds, and that their activity is not specific to any one motor system.

#### 4.4.3 The Role of the Brainstem

One of most prominent differences observed between singing and cello playing was that singing activated the brainstem, including in the PAG, while cello playing did not. The PAG is one of the key regions identified through singing research as instrumental to producing vocalizations (Kleber et al. 2014), a finding which we replicated in the present study. However, the lack of activation in cello playing suggests that not all regions in the vocal-motor network are co-opted in cello playing. While it may be the case that regions like the ACC and aINS, or even the larynx area of motor cortex, can be re-purposed to coordinate activity of motor systems other than those required to produce vocalizations, lower level physiological systems like the brainstem are likely exclusive to vocal control and respiration. Early animal work on the PAG found that it was the first point at which stimulation produced “normal sounding” vocalizations (Jürgens et al. 1979), and later work found that different types of electrical activity in the PAG directly correlated with adduction and abduction of the the laryngial and respiratory musculature during vocalizations in non-human primates (Larson 1991). While systems that coordinate breathing may come into play for more complex instrumental performance aspects like phrasing which are tightly coupled to respiration (Watkins et al. 2012), these systems may not be directly involved in the hand/arm movements required to produce single notes during the investigated task. Without recording muscle activity of the larynx during cello playing, we cannot rule out the possibility that cello playing causes larynx activity. However, we can say conclusively that playing the cello did not incidentally produce vocalization during this task and, consequently, that the descending signals from the brainstem to the musculature were specific to each instrument.

#### 4.4.4 The Role of the Putamen

Another difference observed between singing and cello playing was that singing activated the putamen while cello playing did not, though both cello playing and singing activated the GP. The finding that cello playing and singing both recruit the GP is consistent with previous work in both keyboard playing (eg. (Parsons et al. 2005)) and singing (eg. (Zarate et al. 2008)). However, research in singing has also shown that recruitment of the putamen is specifically linked to expertise, with expert singers recruiting the putamen to a greater extent than novice singers when compensating for, or ignoring, introduced pitch perturbations (Zarate et al. 2008). In this study, neither experts nor novices recruited the putamen when simply singing single notes without an introduced manipulation. The interpretation given to this finding in Zarate (2008) is that the putamen is likely involved in correcting for perceived errors in auditory feedback, and that singing single tones was simple enough that no real error correction was needed. They also note that lesions of the putamen have been linked to dysarthria (Jürgens 2002). Putamen activity has also been linked to over-learned automatic responses in motor sequence learning across a number of studies (Lehéricy et al. 2005; Penhune et al. 2012). In our research, no feedback manipulation was introduced either during cello playing or singing. As such, it could be the case that participants were correcting for incorrect intonation during singing and not cello playing. However, both singing and cello playing show a higher degree of pitch variability at the start of trials, compared to the end. This suggests that corrections to produced pitch were being carried out in both cases. Following from this, it could be the case that the putamen was not recruited for cello playing during our experiment because the function of the putamen is unique to the vocal domain. However, this would be in conflict with findings regarding the putamen’s role in other types of sensorimotor adaptation (eg. (Seidler et al. 2006)) and auditory-motor integration tasks like tapping to the beat (Kung et al. 2012). More likely, it is the case that, similar to the aINS and ACC, the putamen servers a more domain general role in auditory-motor integration, and was not shown to be active for cello playing due to a lack of statistical power. We predict that the putamen will be recruited for both cello playing and singing in future studies involving tasks that more directly probe for auditory-motor integration.

#### 4.4.5 The association between auditory and motor regions

This study also provides direct evidence supporting the idea that playing single notes on the cello not only recruits many of the same brain areas as singing, but that it makes use of the same network of brain regions. First, we found that good intonation is positively correlated with functional brain activity within the areas that are recruited both for singing and cello playing Fig. 9. This shows that the same activity in both instruments plays a role in accomplishing the same behavioural goal. Second, we found that auditory (bilateral HG) and motor (dorsal pre-motor, SMA) regions within the areas common to singing and cello playing were functionally connected during both tasks, and that the degree of functional connectivity is positively correlated with good intonation Fig. 10. In other words, the same brain areas are working together to accomplish both singing and playing the cello in tune during the presented task. This finding addresses the potential criticism that the brain areas observed in the GLM analyses are coincidently active but not necessarily interacting. The functional connectivity findings are also consistent with previous work showing that singing recruits a functionally connected network of brain areas (Zarate et al. 2010)

**Fig 10.**
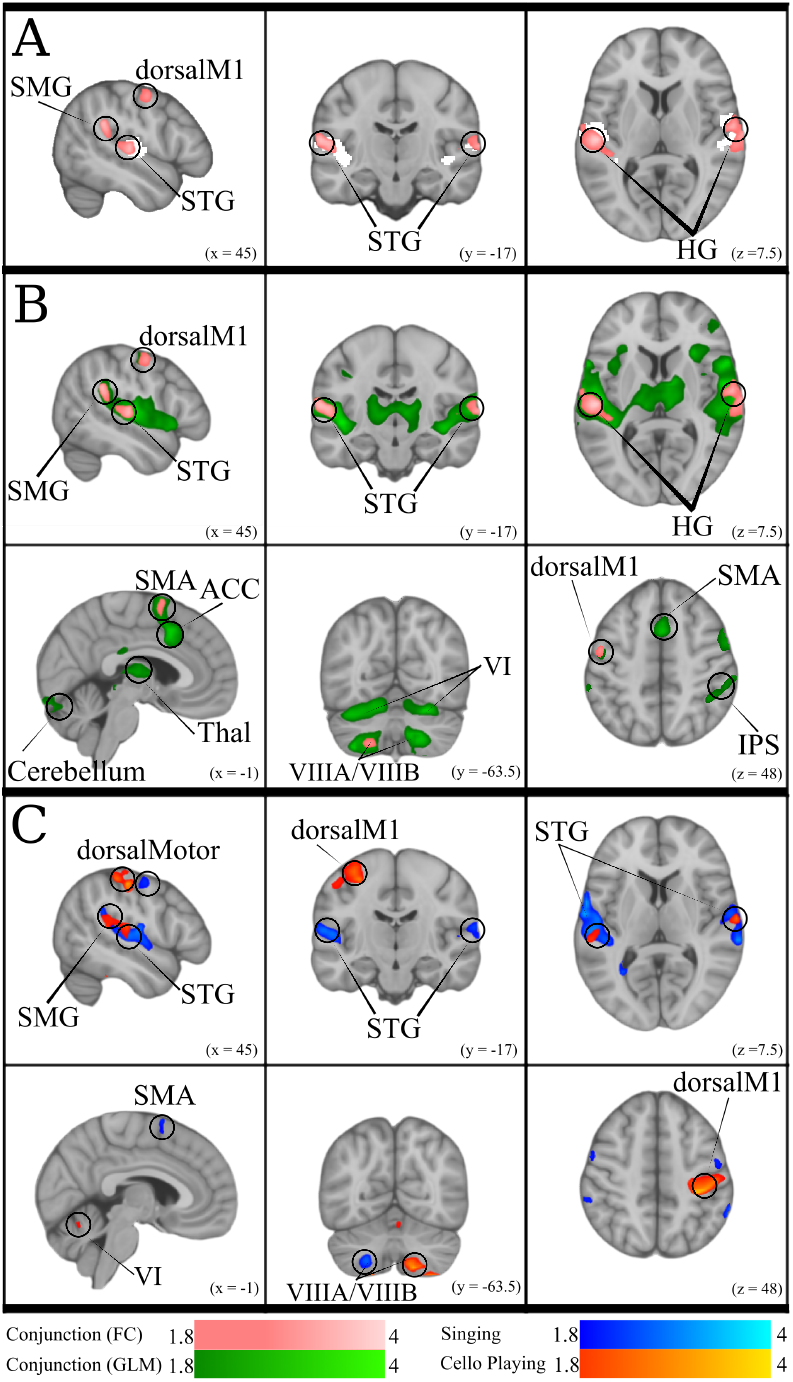
A: Conjunction of functional connectivity during Singing and Cello Playing (pink) overlayed on bilateral Heschle’s Gyrus seed region (white). B: Conjunction of Functional Connectivity during Singing and Cello Playing (pink) overlayed on Conjunction of simple singing and cello playing (green). C: Functional connectivity for Cello Playing > Singing (orange) and Singing > Cello Playing (blue)

### 4.5 Questions Not Answered

#### 4.5.1 STG Recruitment

Even when directly comparing singing and cello playing within the same individuals, some questions are still unanswered. For instance, we found increased activation in cello playing relative to singing in STG. There are two main ways that increased activation may be interpreted when activity falls within regions that are active for both tasks. The first is that increased activation is a sign of enhanced processing. The second is that it is a sign of decreased cognitive efficiency. We propose that the years of explicit training required to play the cello results in enhanced processing during cello tasks relative to singing. This interpretation is consistent with previous work in singing, which has shown that expert singers recruit STG to a greater extent than do novices (Zarate et al. 2008) and previous work in trumpet players showing preferential recruitment of STG during trumpet playing (Gebel et al. 2013). Following from this, we would predict that if expert singers were compared to expert cellists there would be less of a difference in STG activity levels. A third possible interpretation is that the difference is the result of a confounding factor like intensity or another physical feature of the sound.

#### 4.5.2 Single notes vs Melodies

This experiment used a listen/play paradigm with single tones as opposed to melodies, which may limit how well these findings generalize to musical performance in a more naturalistic context. In this regard it may be important to consider our findings in the context of continuous feedback hit-track paradigms. For instance, work done on goal directed movements has characterized an open loop, goal directed and closed loop, feedback oriented system for motor control (for review: (Gaveau et al. 2014)) and brain activity measured using single note reproduction tasks like the one used in this study may be biased towards the open loop component. However, we do not believe that playing a single tone would bear no similarity at all to playing a pattern of tones. Our findings did show activation in line with that of both singing melodies (Kleber et al. 2007) and playing melodies on musical instruments (Lotze et al. 2003). Most importantly, we recently carried out an fMRI study on learning to produce a four-note sequence on the cello which shows very similar auditory-motor activation in dorsal motor and pre-motor areas to those observed here (Wollman et al. 2018: under review). We therefore conclude that the neural systems are similar for production of a single tone as they are for production of a short sequence.

#### 4.5.3 Discrete vs Continuous Pitch Instruments

One of the premises we out forth in our introduction is that cello playing is uniquely similar to the human voice. However, one could argue that the linear arrangement of keys and, consequently, pitches along the length of a keyboard is more related to the monotonic arrangement of pitch along the human vocal cord than the many-to-one mapping of pitches on the cello. We would argue that the differences between signing and keyboard playing are much more compelling, given that motor control over the larynx entails muscular contraction of the vocal cords to different degrees, coordinated with breathing, whereas to play a keyboard requires coordinated action of muscles, joints, limbs, and possibly body posture. Furthermore, with respect to the point about a monotonic mapping, it is possible to play any of the 88 notes on a piano with any of the ten fingers of the two hands. Therefore there is no one-to-one mapping between motor action and sounded pitch; rather, there is always more than one fingering combination to produce the same pitch. We acknowledge that when all strings and all hand positions are used to play the cello it also creates a many-to-one mapping of action to pitch. However, in our study we specifically limited the task to the use of the index finger on one string to maximize the similarity between our cello and singing tasks. By imposing these limitations (string, finger movement) we control for both the many-to-one mapping of location to pitch, and of action to pitch present in everyday cello playing.

Our research cannot directly address the question of how these findings would generalize to discrete pitch instruments like the keyboard or guitar. These instruments do not allow for the online pitch adjustments that are integral to singing or playing continuous pitch instruments in tune. Therefore, the most important differences are likely to emerge in paradigms that exploit this aspect of on-line correction. However, based on the fact that singing and cello playing show such a high degree of overlap in recruited brain areas despite being such physically different tasks, we would speculate that discrete pitch instruments would show a high degree of overlap as well when no online correction is required. This prediction would also be consistent with the large body of research showing that many of the same brain areas are recruited during both piano playing (Parsons et al. 2005) and guitar playing (Vogt et al. 2007) (for review: (Zatorre et al. 2007)).

### 4.6 Future Directions

Using auditory feedback to meaningfully alter movements is one of the core features of auditory-motor integration. In the present study we did not specifically test how auditory feedback affected motor output. As such it is possible, though unlikely, that our participants were relying exclusively on the feed-forward component of the auditory-motor integration network and that auditory feedback was not being used to inform their movements. One of the classic ways of studying the neural correlates of auditory-motor integration is to use pitch perturbation paradigms, where participants are specifically instructed to compensate for introduced perturbations in auditory feedback (Burnett et al. 1998; Zarate et al. 2010)). In so doing, researchers can directly observe which brain regions are involved in integrating auditory feedback with motor planning and execution. Using such paradigms in future experiments will allow us to observe how this auditory-motor integration occurs in cello playing and once again compare these findings with singing.

Another axis along which singing and cello playing might differ, even if the same brain areas are recruited, is the timing of the different processing steps. For instance, it could be the case that the auditory-motor integration network processes and responds to pitch perturbations more quickly during vocal tasks than cello playing due to the evolutionary significance of the voice, and/or due to connectivity differences between auditory and motor systems involved. Directly comparing the latency of event related potentials during both singing and cello playing would allow us to address this question. In doing so, we would gain a more complete understanding of how new skills make use of existing mechanisms in the brain for accomplishing similar tasks.

## Acknowledgments

The authors gratefully acknowledge the work of of Han-Byul Moon for segmenting many audio files, and Luke Zawadiuk for recruiting and scheduling participants, helping collect data, and segmenting many (many) audio files. They also acknowledge Yves Méthot and the MRI technicians for their technical assistance, and Boris Kleber for his support. This work was supported by an operating grant from the Canadian Institutes of Health Research to Robert Zatorre and Virginia Penhune.

